# Exploring the evolutionary divergence of cyclic di-nucleotide signaling in diverse mycobacterial species

**DOI:** 10.1101/2025.09.16.676448

**Authors:** Sayantan Mitra, Sandip Paul, Kamakshi Sureka

## Abstract

Cyclic dinucleotides (CDNs), such as cyclic di-AMP (c-di-AMP) and cyclic di-GMP (c-di-GMP), are key second messengers that regulate fundamental bacterial processes, including cell wall synthesis, biofilm formation, antibiotic resistance, stress response, and virulence. These pathways are particularly relevant in major pathogens like *Mycobacterium tuberculosis*. Increasing evidence highlights the pathogenic potential of non-tuberculous mycobacteria (NTM), originally environmental species that are emerging as significant human pathogens. Understanding the evolution of CDN signaling may therefore provide critical insights into this transition. Our comparative genomic analysis revealed that the c-di-AMP synthase *disA* is present as a single copy in nearly all mycobacterial genomes, except within the genus *Mycolicibacter*. The corresponding phosphodiesterases, PDE and AtaC, are variably distributed, with pathogenic mycobacteria showing a preference for PDE over AtaC. In contrast to the relatively conserved c-di-AMP system, the c-di-GMP pathway comprising diguanylate cyclases (DGCs) and phosphodiesterases (PDEs), exhibits remarkable variation in gene presence/absence and domain architecture across the *Mycobacteriaceae* family. This diversity suggests multiple independent gene gain and loss events throughout evolution, often accompanied by the acquisition of accessory domains. Evolutionary analyses indicate that *disA*, like core housekeeping genes, is under strong purifying selection, while *pde* shows relatively low diversity, reinforcing the idea that c-di-AMP signaling is highly conserved and essential across mycobacteria. By contrast, the c-di-GMP system is more heterogeneous, reflecting its role in environmental adaptation. Together, these findings highlight a sharp evolutionary divergence in CDN signaling, where c-di-AMP appears indispensable for core physiology, whereas c-di-GMP provides flexibility for niche-specific adaptation within the *Mycobacterium* genus.

## INTRODUCTION

The *Mycobacterium* genus encompasses a phylogenetically diverse array of bacterial species that occupy a wide range of ecological niches, including terrestrial and aquatic environments, as well as human and animal hosts (1,2,3,4). This genus includes several medically significant pathogens, most notably *Mycobacterium tuberculosis*, the causative agent of tuberculosis (TB), which remains a major global health threat. Approximately 98% of human TB cases are attributable to *M. tuberculosis*, a member of the *M. tuberculosis* complex (MTBC), which also comprises related species such as *M. bovis*, an animal pathogen capable of zoonotic transmission (5,6). Another prominent species, *M. leprae*, is the etiological agent of leprosy and is characterized by an obligate intracellular lifestyle and a highly reduced genome (7). Beyond the MTBC, the genus includes a large and diverse group of non-tuberculous mycobacteria (NTM), which are primarily environmental saprophytes but have emerged as opportunistic pathogens, particularly in immunocompromised hosts (8,9,10). Among NTMs, the *M. avium* complex (MAC) and *M. abscessus* complex (MABC) are clinically significant due to rising prevalence, multidrug resistance, and links to chronic pulmonary disease (10,11). In response to growing recognition of the phylogenetic and functional heterogeneity within the genus, recent taxonomic revisions based on comprehensive phylogenomic and comparative genomic analyses have proposed a reclassification of the genus *Mycobacterium* into five distinct monophyletic lineages: *Mycolicibacterium, Mycolicibacter, Mycolicibacillus, Mycobacteroides*, and an emended *Mycobacterium* (12).

The wide ecological and pathogenic spectrum observed across the *Mycobacterium* genus suggests that species have evolved diverse physiological and molecular strategies for survival, host interaction, and persistence. Key to these adaptive strategies are small-molecule second messengers, particularly cyclic di-nucleotides (CDNs), which mediate the bacterial response to environmental stimuli and regulate fundamental processes such as cell wall integrity, stress resistance, biofilm formation, immune modulation, and virulence (13,14,15). Among these, cyclic di-AMP (c-di-AMP) and cyclic di-GMP (c-di-GMP) have emerged as central signaling molecules in mycobacterial physiology. In *M. tuberculosis* and *M. smegmatis*, c-di-AMP regulates DNA repair, immune activation, morphology, biofilm suppression, and iron metabolism. (16,17,18). Elevated levels reduce virulence but impair DNA repair and cell wall integrity via effectors like RecA and DarR (19,20,21). c-di-AMP is synthesized by diadenylate cyclases (DACs) and its degradation is mediated by its specific phosphodiesterases (PDEs) maintaining the intracellular homeostasis (22,23,24). Similarly, c-di-GMP plays a crucial role in biofilm development, antibiotic resistance, oxidative stress response, and dormancy regulation (25,26,27). In *M. smegmatis*, it influences cell envelope composition, colony morphology, and motility, while in *M. tuberculosis*, it has been implicated in survival under anaerobic and nutrient-limiting conditions (18,26,27). c-di-GMP synthesis and degradation are governed by diguanylate cyclases (DGCs) and PDEs harboring GGDEF, EAL domains, or a relatively newly described HD-GYP domain, respectively (28). Its regulatory targets include LtmA, HpoR, EthR, DevR, and Lsr2, linking it to diverse transcriptional and metabolic networks (29,30,31).

Despite these insights, the current understanding of CDN-mediated signaling remains largely restricted to a few model and pathogenic species, notably *M. tuberculosis* and *M. smegmatis*. This leaves a substantial knowledge gap regarding the prevalence, diversity, and functional significance of CDN signaling systems in the broader mycobacterial lineage, particularly in emerging NTMs. Comparative genomic and phylogenetic analyses are therefore needed to uncover the conserved and divergent features of CDN pathways across the genus. In this study, we systematically investigate the distribution and evolutionary divergence of genes involved in c-di-AMP and c-di-GMP signaling across a broad spectrum of species within the genera *Mycolicibacterium, Mycolicibacter, Mycolicibacillus, Mycobacteroides*, and an emended *Mycobacterium*. Using comprehensive bioinformatic approaches, we analyzed the presence, conservation, and diversity of CDN synthases and phosphodiesterases, and asses their evolutionary trajectories to infer functional adaptations.

## RESULTS

### Core genome diversification and distribution of c-di-AMP and c-di-GMP signaling genes

The phylogenetic reconstruction from 73 non-recombinant core genes for five genera within the *Mycobacteriaceae* family revealed nearly five distinct clusters corresponding to each genus (Figure 1, 2). The only notable exceptions are the scattered presence of ten *Mycobacterium* species (*M. kyogaense, M. doricum, M. lehmannii, M. neumannii, M. neglectum, M. syngnathidarum, M. aquaticum, M. dioxanotrophicus*) found within the *Mycolicibacterium* cluster, as well as one species (*M. novum*) in *Mycolicibacter* and another one (*M. talmoniae*) in *Mycolicibacillus*. This clustering pattern may arise due to instances of incorrect genus assignments or unidentified divergence patterns, given the recent division of the former *Mycobacterium* genus into these five new genera (12, 32). In the genus *Mycobacterium*, the MTBC members (*M. tuberculosis, M. africanum, M. bovis, M. orygis, M. microti, M. canettii*) tend to cluster together (Figure 1, 2). This clustering reflects a robust phylogenetic analysis based on non-recombinant core genes. In case of MAC, the component organisms, *M. avium, M. timonense, M. bouchedurhonense, M. arosiense, M. colombiense, M. vulneris, M. marseillense, M. intracellulare, M. intracellulare* subsp. *chimaera, M. intracellulare* subsp. *yongonense*, and *M. paraintracellulare* cluster together with *M. mantenii* (33). This clustering suggests that *M. mantenii* is also part of the MAC. For MABC, belonging to the *Mycobacteroides* genus, the members *M. abscessus, M. chelonae, M. franklinii, M. imunogenum, M. salmoniphilum, M. saopaulense* and *M. stephanolepidis* cluster together separately (34,35,36,37).

**Figure 1.**
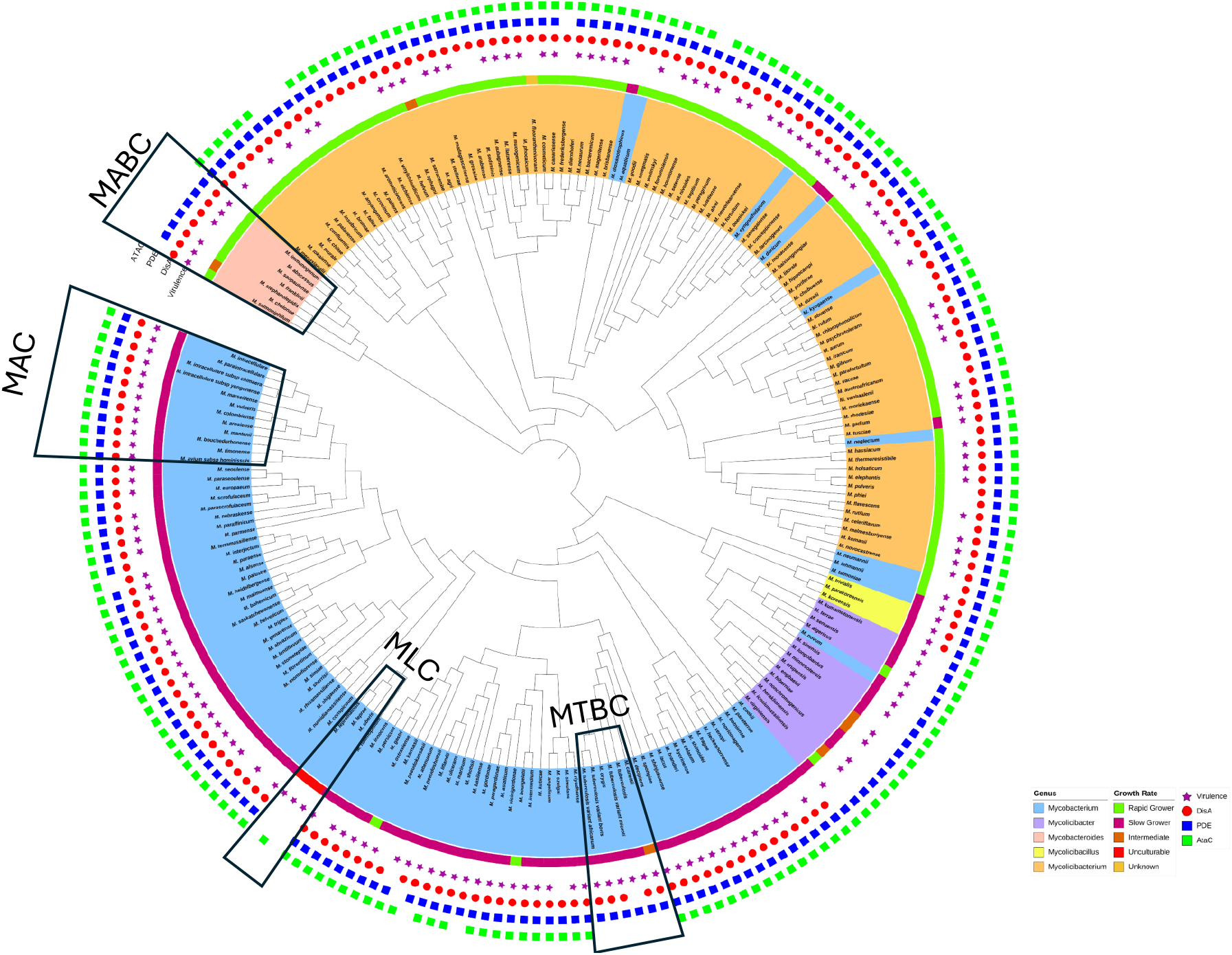
Core genome phylogeny along with depiction of distribution of c-di-AMP signaling pathway genes *disA* (red circle), *pde* (blue square), and *ataC* (green square) an Virulence (star)..

**Figure 2.**
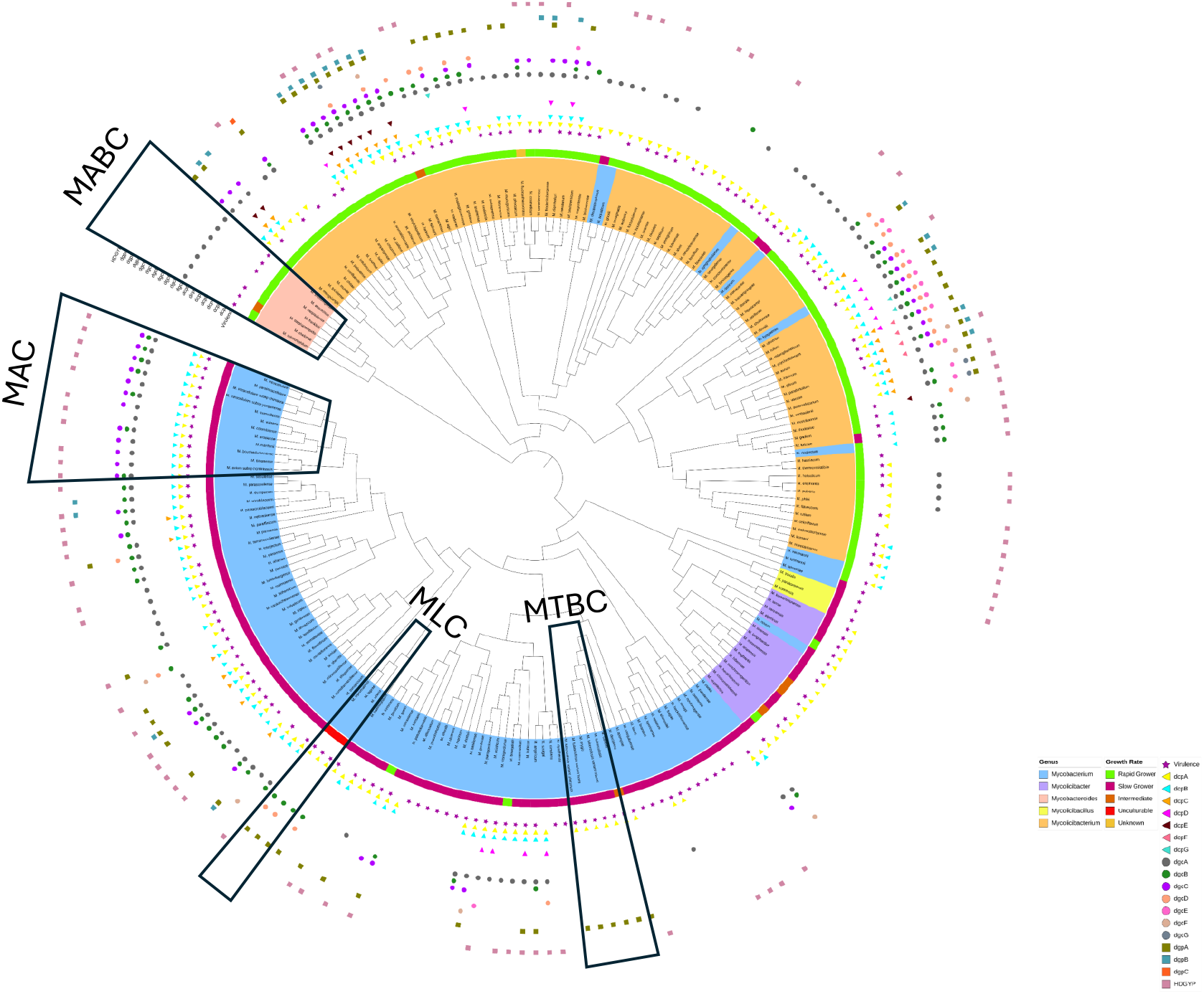
Core genome phylogeny along with depiction of distribution of c-di-GMP signaling pathway genes *dcp*A-G, *dgc*A-G, *dgp*A-C, HDGYP and Virulence (star).

In the c-di-AMP signaling pathway, DisA and two c-di-AMP-specific phosphodiesterases (PDE and AtaC) were searched across 214 *Mycobacteriaceae* genomes and the results are mapped in the core genome phylogeny (Figure 1). These genes are found to be widely distributed, with *disA, pde*, and *ataC* present in 90.65%, 96.73%, and 90.19% of genomes, respectively. Within the MTBC group, *ataC* is consistently absent, whereas *disA* and *pde* were retained, a pattern also observed in the MABC group, suggesting a genus-specific feature *of Mycobacteroides*. In contrast, the *disA* gene is completely absent in *Mycolicibacter* genus, including *M. novum*, which clusters within this genus based on core genome phylogeny. Furthermore, truncated form of *disA* is present in the reduced genomes (*M. leprae, M. lepromatosis* and *M. uberis*) and absent from the genomes *M. tuberculosis* variant *microti* and *M. pseudoshottsii* within the genus *Mycobacterium*. Overall, these findings indicate that while c-di-AMP signaling proteins are broadly conserved in *Mycobacteriaceae*, specific gene losses characterize certain lineages (Figure 1).

The intracellular homeostasis of c-di-GMP is regulated by two opposing enzymatic activities: diguanylate cyclases (DGCs), which synthesizes c-di-GMP from two molecules of GTP, and phosphodiesterase (PDEs) which degrade it to pGpG (28). DGCs typically harbor a catalytically active GGDEF domain, often preceded by regulatory domains such as GAF or PAS, whereas PDEs contain either an EAL domain or an HD-GYP domain. Additionally, many hybrid proteins have been identified that harbor both GGDEF and EAL domains. In *Mycobacteriaceae*, we have identified a diverse set of c-di-GMP-related proteins with varied domain architecture. Notably, Sharma *et al*. renamed MSDGC-1 from *M. smegmatis* as DcpA (<u>d</u>iguanylate <u>c</u>yclase and <u>p</u>hosphodiesterase A) due to its GAF-GGDEF-EAL configuration (38). Following this convention, we have designated names to all hybrid GGDEF-EAL as DcpB through DcpG, ordered by their abundance across genomes. Proteins with only a diguanylate cyclase domain, either alone or in combination with GAF, PAS, or PAC domains, have been named DgcA to DgcF. Standalone PDEs, either alone or in combination with GAF or PAS domains, have been named DgpA to DgpC. All c-di-GMP signaling genes were screened across 214 genomes and mapped to the core genome phylogeny (Figure 2, Table 1).

**Table 1.** Names of the various DGCs and PDEs according to their domain organization and abundance in Mycobacteriaceae family

| Domain combination | # of representative organisms | Assigned name | Type of protein |
| --- | --- | --- | --- |
| **GAF-GGDEF-EAL** | 171 | *dcpA* | Hybrid |
| **GGDEF-EAL** | 108 | *dcpB* | Hybrid |
| **GAF-PAS-PAC-GGDEF-EAL** | 24 | *dcpC* | Hybrid |
| **PAS-PAC-GGDEF-EAL** | 14 | *dcpD* | Hybrid |
| **REC-GAF-PAS-GGDEF-EAL** | 9 | *dcpE* | Hybrid |
| **REC-GAF-PAS-PAC-GGDEF-EAL** | 4 | *dcpF* | Hybrid |
| **GAF-PAS-GGDEF-EAL** | 3 | *dcpG* | Hybrid |
| **GGDEF** | 129 | *dgcA* | DGC |
| **PAS-PAC-GGDEF** | 70 | *dgcB* | DGC |
| **GAF-GGDEF** | 53 | *dgcC* | DGC |
| **PAS-GGDEF** | 27 | *dgcD* | DGC |
| **GAF-PAS-PAC-GGDEF** | 13 | *dgcE* | DGC |
| **GAF-PAS-GGDEF** | 8 | *dgcF* | DGC |
| **PAC-GGDEF** | 3 | *dgcG* | PDE |
| **EAL** | 57 | *dgpA* | PDE |
| **GAF-EAL** | 23 | *dgpB* | PDE |
| **PAS-EAL** | 1 | *dgpC* | PDE |
| **HD-GYP** | 85 |  | PDE |

Hybrid proteins are particularly widespread, with DcpA (GAF-GGDEF-EAL) being the most prevalent (79.91% of genomes). Other abundant proteins included DgcA (GGDEF-only, 60.28%), DcpB (GGDEF-EAL, 50.46%), HD-GYP domain proteins (39.72%), DgcB (PAS-PAC-GGDEF, 32.71%) and DgpA (EAL-only, 26.64%). Additional hybrids (e.g., DcpC, DcpD, DcpE, DcpF and DcpG) and proteins with single domains (GGDEF/EAL) or with different accessory domains (such as DgcC, DgcD, DgcE, DgcF, DgpB and DgpC) are also detected (Figure 2).

The distribution of these genes was highly uneven across the family. The genus *Mycolicibacillus* lacks any c-di-GMP signaling genes, while in *Mycolicibacter*, only *dcpA* is consistently present. In *Mycobacteroides*, only *dgcA* gene is present for 7 members. Furthermore, there are 4 genomes in the genus *Mycolicibacterium* and 14 in the genus *Mycobacterium* including a cluster of *M. ulcerans* and other five closely related genomes belonging to the same clade that completely lack c-di-GMP pathway genes (Figure 2). Within the *Mycobacterium* genus, gene repertoire clustered into two groups: *M. leprae* and its closely associated genomes, and *M. kubicae* along with its related genomes. In *Mycolicibacterium*, three clusters are present: *M. anyangense, M. madagascariense*, and *M. hippocampi* and their respective relatives.

Together, these results indicate that c-di-AMP signaling is more universally conserved across *Mycobacteriaceae* than c-di-GMP signaling, which shows a patchy distribution. The diverse repertoire of c-di-GMP genes, shaped by domain composition and lineage-specific gene retention or loss, reflects its specialized and non-uniform role in mycobacterial physiology.

### Diversification of c-di-AMP signaling pathway genes and functional domains

Phylogenetic and domain architecture analyses revealed distinct patterns of divergence among c-di-AMP signaling proteins in *Mycobacteriaceae*. DisA, which is present as a single copy in the genomes of the *Mycobacteriaceae* family, is known to have the most complex domain architecture among all known DACs. In the protein sequence phylogeny, proteins from the genera *Mycobacterium* and *Mycolicibacillus* cluster separately, while proteins from the genus *Mycobacteroides* form a distinct cluster within *Mycolicibacterium*, indicating close sequence similarities between these two genera for the DisA proteins (Figure 3). The domain architecture of nearly all proteins follows a similar pattern, featuring an N-terminal DisA_N domain and a C-terminal helix-hairpin-helix (HhH) domain, connected by a long DisA-linker domain, as previously noted (39). However, there is an exception where the HhH domain is absent in six genomes (*M. heckeshornense, M. xenopi, M. numidiamassiliense, M. malmoense, M. saskatchewanense*, and *M. shottsii*) within the genus *Mycobacterium* (Figure 3). Overall, the domain architecture of the DisA protein is conserved within this family.

**Figure 3.**
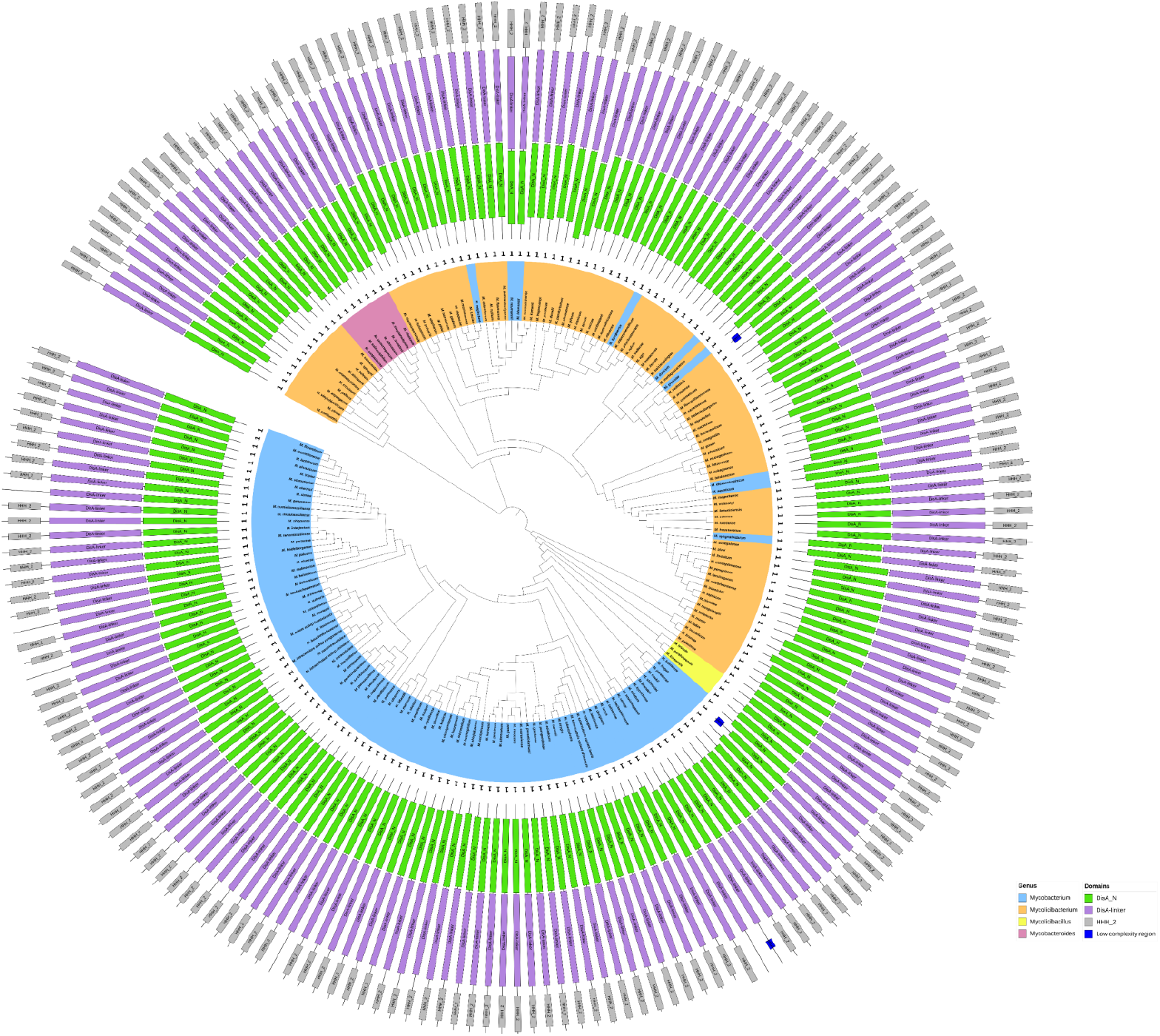
Phylogeny of c-di-AMP Signaling Pathway Gene *dis*A along with depiction of distribution of functional domains DisA_N, DisA-linker, HHH_2, low complexity region

The most prevalent phosphodiesterase protein (PDE) is present as a single copy in all five genera and consists of the conserved DHH and DHHA1 domains, in *M. flavescens*, only the DHHA1 domain is present which could be an inactive form of PDE as the DHH domain is known to be essential for phosphodiesterase activity (Supplementary Figure 1) (22). The phylogeny of this PDE protein closely mirrors that of the core genome, although it is absent from seven genomes (*M. leprae, M. lepromatosis, M. uberis, M. liflandii, M. interjectum, M. frederiksbergense* and *M. salmoniphilum*). This suggests that the diversification of the PDE protein followed the pattern of core gene diversification, with some organisms losing the protein during genome evolution. Many of the PDE proteins exhibit small predicted low-complexity regions, and their consistent presence in conserved locations across specific genome clusters indicates a similar pattern of genomic divergence within these groups.

The phosphodiesterase protein AtaC is absent in the members of the genus *Mycobacteroides*. This protein clusters separately for the genomes of the genus *Mycolicibacter* probably due to independent events of this gene gain (Supplementary Figure 2). Most AtaC proteins contain a single phosphodiesterase domain, but some proteins in certain *Mycolicibacter* and *Mycolicibacterium* genomes also feature a signal peptide and transmembrane domain. Proteins of these genomes tend to cluster together. Notably, many of these organisms possess multiple copies of the AtaC protein, whereas in the genera *Mycobacterium* and *Mycolicibacterium*, the protein is typically present as a single copy.

### Diversification of c-di-GMP signaling pathway genes and functional domains

The genomes of the *Mycobacteriaceae* family encode a diverse repertoire of enzymes regulating c-di-GMP synthesis and degradation, classified into 18 types based on conserved domains (GGDEF, EAL, HD-GYP) and associated accessory domains (PAS, GAF, REC, PAC). Additional features such as transmembrane segments and coiled-coil regions are also mapped onto protein sequence phylogeny. The most widespread hybrid protein, DcpA (GAF-GGDEF-EAL), is detected across the genera *Mycobacterium, Mycolicibacterium*, and *Mycolicibacter* (Figure 4). This protein shows a conserved domain architecture. DcpB is restricted to *Mycobacterium* and *Mycolicibacterium* and contains both GGDEF and EAL domains, along with transmembrane domains, indicating that it functions as an integral membrane protein (supplementary figure 3a). Most proteins in the *Mycolicibacterium* possess a single GGDEF domains, whereas in *Mycobacterium* DcpB proteins poses a pair of GGDEF domain. Also copies of *dcpB* gene up to 7 can be seen to be present in five of the genomes of the genus *Mycolicibacterium*. The two other hybrid proteins DcpC and DcpD (Supplementary figures 3b and 3c) are present exclusively in the genera *Mycobacterium* and *Mycolicibacterium* same as DcpB, while three others (dcpE, dcpF and dcpG, Supplementary figure 3e-g) are present in few organisms exclusively in the genus *Mycolicibacterium*.

**Figure 4.**
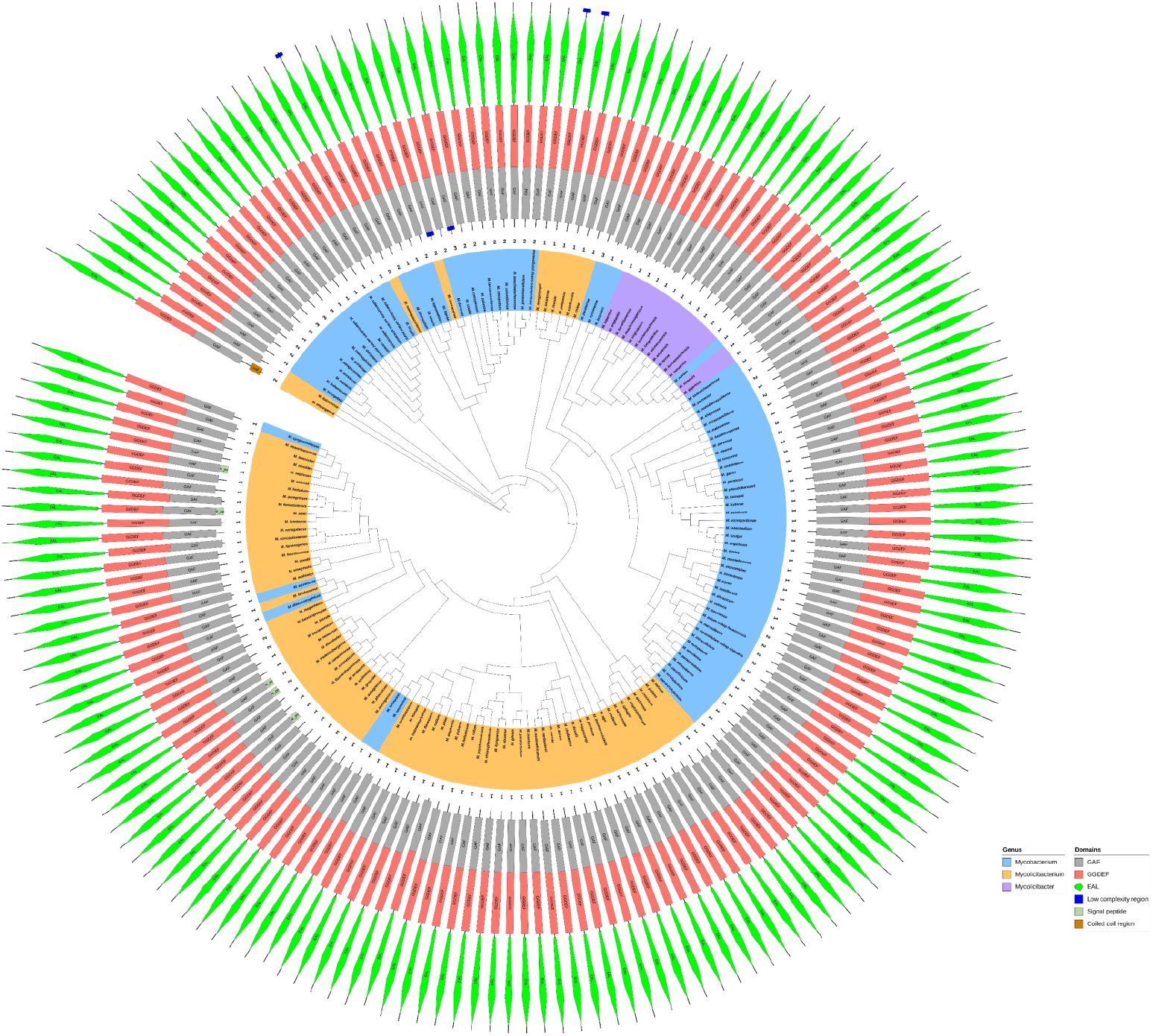
Phylogeny of c-di-GMP Signaling Pathway Gene *dcp*A along with depiction of distribution of functional domains GAF, GGDEF, EAL, low complexity region, signal peptide, coiled coil region

Among diguanylate cyclases, DgcA (GGDEF-only) is the most abundant, present in *Mycolicibacterium, Mycobacterium*, and *Mycobacteroides*. In MABC group, *dgcA* is the sole c-di-GMP signaling gene identified, typically associated with transmembrane and sometimes MASE1 domains (Supplementary Figure 4). Other GGDEF proteins (DgcB–DgcF) with combinations of GAF, PAC, or PAS domains were restricted to *Mycobacterium* and *Mycolicibacterium*, while DgcG (GGDEF–PAC) was found only in three *Mycolicibacterium* species (Supplementary figure 5a-f).

There are three combinations of phosphodiesterase proteins based on single EAL domain or EAL with GAF/PAS domain found in the *Mycobacterium* and *Mycolicibacterium* genera. They are: DgpA (EAL-only), DgpB (GAF-EAL), and DgpC (PAS-EAL). Among these, DgpA is the most widely distributed, found predominantly in the *Mycobacterium* and *Mycolicibacterium* genera (Supplementary Figure 6a). Notably, only one *dgpA* gene, identified in *M. psychrotolerans*, contains a transmembrane domain. It is also worth mentioning that nearly all members of the MTBC group carry both *dgpA* and *dcpA*. The other two EAL domain-containing genes, *dgpB* is exclusively present in the *Mycobacterium* and *Mycolicibacterium* genera, whereas *dgpC* is present only in *M. palauense* (Supplementary Figure 6).

The second most prevalent protein type is HD-GYP family, present in the *Mycobacterium* and *Mycolicibacterium* genera (supplementary figure 7). These multidomain proteins carried conserved HDc and helix-turn-helix DNA-binding domains, frequently included low-complexity regions, and in some cases contained a signal peptide, particularly in *M. shimoidei, M. mantenii, M. parascrofulaceum, M. scrofulaceum, M. seoulense* and *M. paraseoulense*.

### Diversity analysis of core genome and signaling pathway genes

Among the 73 core genes, the pairwise nucleotide diversity (π) ranged from 0.072 to 0.214, with a mean of 0.153. Except for *disA*, all cyclic-di-nucleotide signaling pathway genes exhibited π values exceeding the upper range observed in the core genes (Figure 5a). Correspondingly, Z-score analysis revealed significant (p<0.05) deviations for all signaling genes except *disA*. A similar trend was observed for the nonsynonymous substitution rate (dN), where core genes displayed a mean value of 0.065 (range: 0.005–0.129) (Figure 5c). For the synonymous substitution rate (dS), *disA, pde*, and *dcpF* had values within the range observed for core genes (mean dS: 0.668, range: 0.363–0.999) and showed non-significant Z-scores (Figure 5b). Notably, *dcpF*, the only c-di-GMP–related gene in this dataset, was detected in just four genomes of the genus *Mycolicibacterium*.

**Figure 5.**
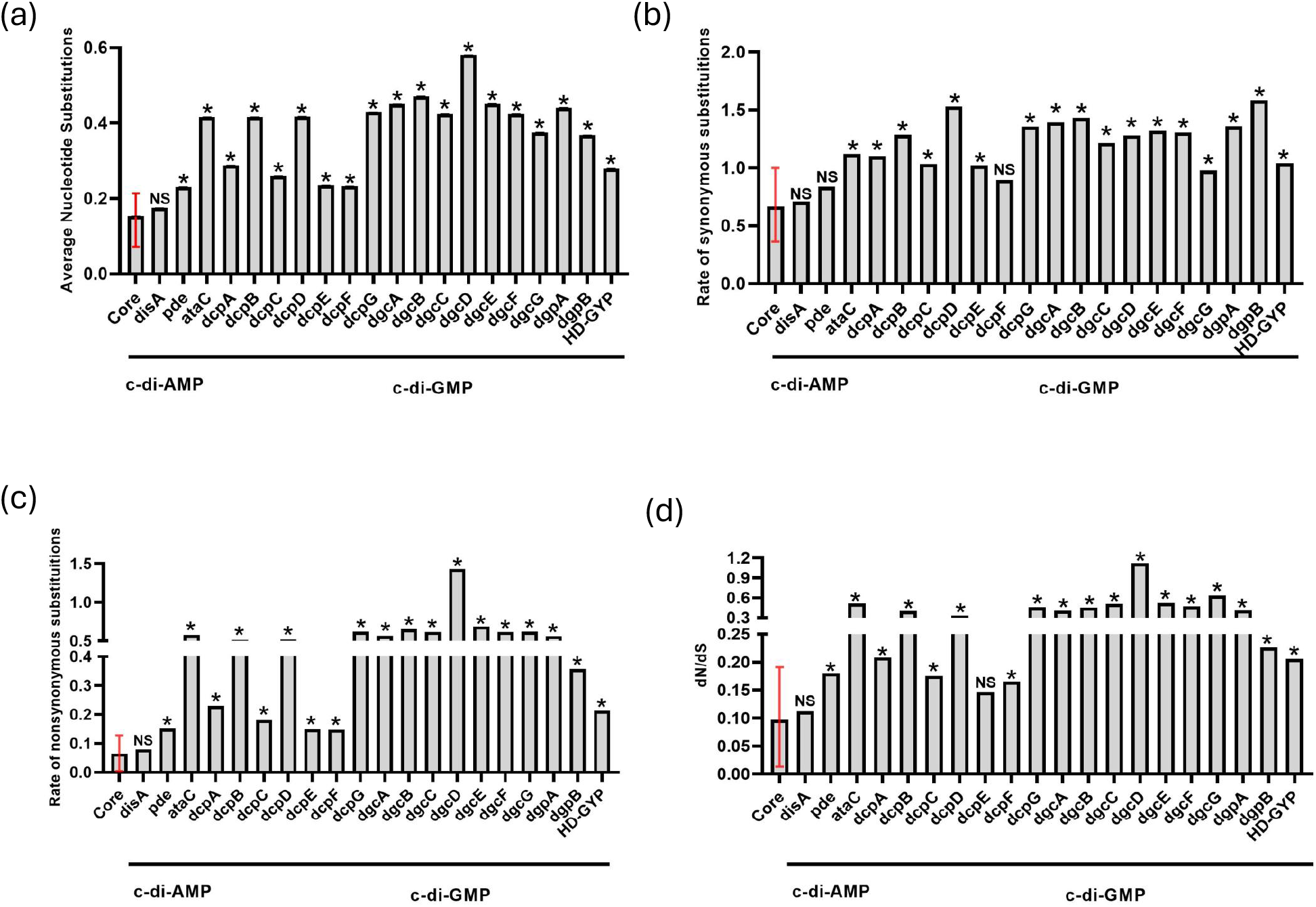
(a, b, c, d): Bar plot depicting average nucleotide diversity, rate of synonymous substitutions (dS), rate of non-synonymous substitutions (dN) and dN/dS for core genes and c-di-AMP and c-di-GMP signaling pathway genes respectively. “^*^” indicates statistically significant deviations (p <0.05, one tailed), and “NS” denotes non-significant deviations, relative to core gene values, as determined using the z-score test.

Core genes were under strong purifying selection, as evidenced by a markedly lower average rate of nonsynonymous substitutions compared to synonymous substitutions (mean dN/dS = 0.097, range: 0.013–0.191). This pattern reflects strong selective pressure against amino acid changing mutations, consistent with the conserved functional roles of housekeeping genes. The comparison of the dN and dS values for each core gene (P < 0.05) showed that dN/dS was significantly less than 1 for all 73 core genes, indicating a predominant occurrence of synonymous changes. In contrast, most cyclic di-nucleotide signaling pathway genes exhibited significantly higher dN/dS ratios (Z-score test, p<0.05) compared to the core genes distribution, although their values still significantly lower than 1, indicating that they remain under overall purifying selection (Figure 5d). Only *disA* and *dcpE* did not differ significantly from the core gene pattern; notably, *dcpE* was identified in only nine genomes within the genus *Mycolicibacterium*. Strikingly, *dgcD*, detected in 27 genomes across the genera *Mycobacterium* and *Mycolicibacterium*, was the only gene found to be under positive selection, with a dN value significantly exceeding the dS value (dN/dS > 1, P < 0.05).

## DISCUSSION

Cyclic dinucleotides (CDNs), such as cyclic di-AMP (c-di-AMP) and cyclic di-GMP (c-di-GMP), serve as essential second messengers in bacteria, regulating a wide range of cellular processes. Our comparative genomic analysis across five genera of the *Mycobacteriaceae* family highlights both the conserved and divergent evolutionary trajectories of these signaling systems. Among CDNs, c-di-AMP occupies a unique position as an “essential poison”: indispensable for bacterial survival but toxic when over-accumulated (40). In *Mycobacterium*, c-di-AMP is not strictly essential in vitro, yet it plays critical roles in physiology and infection, with dysregulation leading to impaired survival during host colonization (16, 41, 42). Its homeostasis is mediated by DisA and two phosphodiesterases (PDE and AtaC), which are found widely distributed across pathogenic (90.5%) and non-pathogenic (91.3%) species. The consistent presence of DisA (except in *Mycolicibacter*) and PDE genes suggests that c-di-AMP signaling is fundamental and crucial to mycobacterial biology. The absence of DisA in *Mycolicibacter*, a genus belonging to the Terrae clade with distinct lipid and cell wall architecture, may reflect adaptive gene loss linked to altered cell wall metabolism (43). Similarly, *M. microti* lacks DisA and displays attenuated virulence, consistent with its historical use as a vaccine candidate (44,45).

Obligate intracellular mycobacteria carry only truncated c-di-AMP genes, likely reflecting adaptation for long-term immune evasion, as c-di-AMP triggers host type I interferon responses. In *M. leprae*, this loss may also relate to its reliance on host lipid metabolism, with many lipid-associated genes reduced to pseudogenes (7). Most *Mycobacteriaceae* encode two PDE types (DHH-DHHA1 and AtaC), ensuring stringent control of c-di-AMP, though MTBC and MABC genomes typically retain only a single *pde* copy and lack *ataC*. The consistent presence of single *disA* and *pde* genes in pathogenic species suggests a link between minimal c-di-AMP machinery and human pathogenicity.

The widespread presence of DisA may be attributed to its dual role in the DNA damage response. However, some members of the *Mycobacteriaceae* family lack the C-terminal DNA-binding domain while retaining the N-terminal DAC domain. This highlights the importance of c-di-AMP signaling, as experimental evidence shows that DisA retains DAC activity even in the absence of the C-terminal DNA-binding domain (46). Phosphodiesterases such as PDE and AtaC are also broadly conserved, and the simultaneous absence of both is rare. PDEs show conserved architectures, though a distinct subset of AtaC proteins carrying additional N-terminal 22aa transmembrane domains suggests potential specialization as membrane-bound PDEs (Supplementary Figure 2). This cluster comprises 19 species with multiple copies of AtaC protein (Except for *M. sarraceniae*), including five from the genus *Mycolicibacter*, four from *Mycolicibacterium*, and one species (*Mycobacterium kyogaense*) from the *Mycobacterium* genus, all of which contain the transmembrane domain. The presence of these transmembrane domain–containing AtaC variants warrants further investigation, particularly into their potential role as membrane-bound phosphodiesterases, similar to PgpH (47).

In contrast to the relatively conserved c-di-AMP system, the c-di-GMP signaling pathway exhibits remarkable variability in both gene copy number and domain composition across the *Mycobacteriaceae*. Turnover proteins are abundant in *Mycobacterium* and *Mycolicibacterium*, entirely absent in *Mycolicibacillus*, and restricted to a single copy in *Mycolicibacter* and *Mycobacteroides*. This heterogeneity, often coupled with the presence of multiple accessory domains, indicates that repeated, independent gene gain and loss events have shaped the evolution of this pathway. Across genomes, the abundance of c-di-GMP turnover genes is significantly positively correlated with both genome size and total gene count, while showing a negative correlation with pseudogene content (Supplementary Fig. 8a–d). The variation in c-di-GMP signaling gene distribution appears to be influenced by habitat type and pathogenic potential. Specifically, environmental genomes, with an average size of ∼5.82 Mb, harbour larger genomes than host-adapted genomes (∼4.95 Mb). Similarly, virulent genomes (∼5.67 Mb) tend to have smaller sizes compared to non-pathogenic genomes (∼6.08 Mb). Consistent with previous findings, bacteria with strong host associations generally maintain fewer GGDEF- and EAL-domain proteins, reflecting reduced reliance on complex signaling in stable or persistent pathogenic niches (48). Conversely, species exposed to dynamic or fluctuating environments retain larger repertoires of c-di-GMP enzymes, enabling versatile responses to extracellular cues. Notably, pathogens with multi-host lifestyles, such as *Yersinia pestis* and *Legionella pneumophila*, encode elevated numbers of c-di-GMP turnover genes, consistent with their need to alternate between pathogenic and non-pathogenic states across diverse host environments (49, 50). By contrast, host-adapted pathogens such as *M. tuberculosis* and *M. leprae* show substantial gene loss, consistent with their relatively stable intracellular environments where complex signaling is less critical. The degradation of c-di-GMP pathways in *M. abscessus* remains puzzling but may be linked to its extensive genome remodeling during host adaptation (51).

Functionally, c-di-GMP proteins display high modularity, often integrating GGDEF (synthase) and EAL (phosphodiesterase) domains with sensory domains such as GAF, PAS, or PAC. DcpA, a hybrid GAF–GGDEF–EALprotein, is the most conserved across *Mycobacteriaceae*. The presence of these domains implies signal-dependent, post-translational regulation of c-di-GMP metabolism (52). Despite their shared catalytic roles, the function of each GGDEF and EAL domain-containing protein may be highly context-dependent. The presence of multiple copies raises important questions regarding their functional specialization and spatio-temporal regulation. Exploring the complex regulation, activity coordination, and evolutionary differentiation of c-di-GMP turnover proteins represents an exciting avenue for future research.

We identified 104 core genes conserved across 214 *Mycobacteriaceae* genomes using a stringent 70% sequence identity and alignment length cutoff. In comparison, a previous analysis of 150 genomes with a relaxed 50% threshold reported 1,941 soft-core proteins present in ≥80% of genomes (12). Our stricter approach was aimed at delineating a highly conserved set of core proteins that are universally shared across the family, thereby providing a robust foundation for evolutionary analyses and deeper insights into the fundamental features of core gene evolution within *Mycobacteriaceae*. Analysis of these core genes revealed an average nucleotide diversity (π) of 0.153. This level of diversity is comparable to that reported for the genus *Streptomyces*, which exhibited an average nucleotide diversity of 0.1373 across 44 species and 136 isolates (53). Interestingly, the five genera of *Mycobacteriaceae*, which were historically grouped under the single genus *Mycobacterium*, displayed genus-specific nucleotide diversity values as follows: *Mycobacterium* (π = 0.134; range 0.058–0.183; 102 species), *Mycobacteroides* (π = 0.073; range 0.015–0.166; 7 species), *Mycolicibacillus* (π = 0.100; range 0.042–0.153; 3 species), *Mycolicibacter* (π = 0.077; range 0.028–0.144; 14 species), and *Mycolicibacterium* (π = 0.139; range 0.064–0.196; 88 species). Notably, the recently redefined genera *Mycobacterium* and *Mycolicibacterium*, both represented by a large number of species, exhibited average nucleotide diversity levels comparable to those of *Streptomyces*. In contrast, the remaining three genera showed much lower diversity, reflecting the presence of more closely related species within each group. Furthermore, we found that all core genes, presumed to be essential in nature, are under strong purifying selection. This observation is consistent with previous studies reporting that core genomes are typically shaped by strong purifying selection (54,55,56). Since purifying selection constrains evolutionary rates, essential genes are expected to display higher levels of conservation (57), which is also evident in our dataset. Within the cyclic dinucleotide pathways, we observed that *disA*, a key component of the c-di-AMP pathway, exhibited an average nucleotide diversity well within the range of the identified core genes, with dS, dN, and dN/dS values indicating strong purifying selection like core genes. These features point toward the probable essential nature of the *disA* gene in this group of organisms. Similarly, the abundant gene *pde* displayed relatively low diversity, with nucleotide diversity and substitution rate values, particularly dS, closely resembling those of the core genes. Together, these findings indicate that the c-di-AMP pathway, represented by its central genes *disA* and *pde*, is highly conserved across all five genera. While *disA* shows features characteristic of essential genes, the broader c-di-GMP pathway appears more diverse, as reflected in the variable presence or absence of its component genes and the heterogeneity of their functional domain architectures.

The differential conservation of these signaling pathways highlights a major evolutionary divergence in second messenger utilization across the *Mycobacteriaceae* family. Whereas c-di-AMP appears indispensable and closely tied to core physiological processes, c-di-GMP likely plays a more specialized role in environmental adaptation. The distinct presence/absence patterns of these genes could thus serve as molecular signatures for classifying *Mycobacterium* species and predicting their ecological strategies or pathogenic potential, while also raising questions about potential redundancy or crosstalk between nucleotide signaling systems in species that encode both. Future research should aim to experimentally validate the functionality of these signaling genes in representative taxa with atypical complements and to investigate the regulatory networks downstream of c-di-AMP and c-di-GMP, which may reveal how these pathways shape mycobacterial physiology, stress adaptation, and virulence.

## METHODS

### Organism selection and sequence retrieval

A total of 214 genomes representing five genera within the *Mycobacteriaceae* family, *Mycobacterium*-102, *Mycobacteroides-*7, *Mycolicibacter-*14, *Mycolicibacillus-*3, and *Mycolicibacterium-*88 were downloaded from NCBI’s RefSeq database (58). We selected the high-quality genomes based on chromosomal completeness. A total of 209 species were considered, each represented by a single genome, with the exceptions of *M. tuberculosis*, which included four representative genomes (*M. tuberculosis* H37Rv, *M. tuberculosis_subsp*.*_africanum, M. tuberculosis subsp. bovis*, and *M. tuberculosis subsp. microti*), and *M. intracellulare*, which included three representative genomes (*M. intracellulare, intracellulare subsp. chimaera*, and *M. intracellulare subsp. yongonense*), due to the availability of high-quality genomes. All relevant fna, ffn, and gff files for these genomes were downloaded from NCBI for subsequent analysis.

### Core genome determination and reconstruction of core genome-based phylogeny

The core genome was determined by first constructing a pan-genomic profile using all annotated protein-coding genes (total 11,09,795) from the 214 genomes under investigation. We employed the CD-HIT Suite (59) to cluster all amino acid sequences, applying a threshold of 70% sequence identity and alignment coverage for orthologous clustering. From these clusters, we generated a pan-matrix that depicted the presence and absence of genes across all organisms studied. This pan-matrix was then used to extract 104 core gene clusters, which are present in 100% of the genomes. To ensure core gene uniqueness, only one representative sequence per organism was retained, removing all paralogs and duplicates. Nucleotide sequences of core clusters were then extracted for recombination and evolutionary analyses.

The Recombination Detection Program (RDP5) was utilized to identify potential recombination events in the core genes (60). This software incorporates a variety of recombination detection algorithms based on three different methodologies: phylogenetic (BOOTSCAN, RDP, and SISCAN), substitution (GENECONV, MAXCHI, CHIMAERA, and LARD), and distance comparison (PHYLPRO). We designated a gene region as tentatively affected by a homologous recombination event if it yielded significant results (P < 0.05) in four or more of the recombination detection algorithms. This analysis led to the identification of 31 gene clusters with probable recombination events, which we subsequently excluded, resulting in 73 core gene clusters for phylogenetic analysis. Next, we aligned the amino acid sequences for each cluster using the MUSCLE (61), followed by concatenating the alignments. The resulting concatenated file was then used to construct a maximum likelihood (ML) tree with the JTT (Jones-Taylor-Thornton) model, utilizing 300 bootstrap replicates, implemented in MEGA11 (62).

### Characterization of c-di-AMP and c-di-GMP signaling pathway genes in the genomes

Using the DisA (NP_218103.1) and PDE (NP_217353.1) proteins of c-di-AMP pathway and GGDEF and/or EAL-domain containing proteins (NP_215870.1, NP_215873.1) of c-di-GMP pathway from *M. tuberculosis* as reference sequences, we employed standalone BLASTP to identify the orthologous partners of these genes in the other genomes under study. Since the *ataC* gene is absent in *M. tuberculosis*, we used the AtaC (WP_003894937.1) protein from *M. smegmatis* as a reference. We applied a 50% sequence similarity and length coverage threshold for cyclic di-AMP genes (except for *ataC*, where 20% sequence similarity was used) and a 20% threshold for cyclic di-GMP genes. All BLAST hits were assessed for the presence of full-length genes in each organism, and previous annotations were also reviewed. This process allowed us to characterize both the presence/absence and copy numbers of all signaling pathway genes. Additionally, we used BLASTN to compare reference sequences against genomic fna files, applying the specified thresholds for sequence identity and length coverage to identify any pseudogenic or truncated forms of these genes in the genomes under investigation (Only for C-di-AMP pathway genes).

We further examined all full-length protein sequences identified from the BLASTP analysis for all organisms in the study to check for the presence of conserved domains. This was done using the SMART web resource, with options for PFAM, signal peptides, and internal repeats enabled (63).

### Molecular evolutionary analysis of core genes and signaling pathway genes

For the molecular evolutionary analysis, we conducted DNA codon sequence alignment using PAL2NAL (64), deriving alignments from multiple amino acid sequences and their corresponding nucleotide sequences for 73 core genes and all CDN signaling pathway genes. We calculated average pairwise nucleotide diversity value (π) and subsequently estimated the rates of nonsynonymous (dN) and synonymous (dS) mutations using the mutation-fraction method (65), implemented in TimeZone (66). Z-scores and p-values were computed based on the distribution of core gene values to identify statistically significant deviations among the CDN signaling pathway genes.

To evaluate the significance of differences between dN and dS, we applied a nonparametric bootstrap approach (67), simulating 1,000 datasets by resampling DNA sites from the multiple sequence alignment. This method effectively disrupted the codon structure of the original dataset. Consequently, we generated the distribution of dN/dS under the null hypothesis of neutrality (dN/dS = 1). By comparing the rank of the observed dN/dS value within the simulated distributions, we were able to identify the presence of positive selection, negative selection, or neutrality, at 95% significance level.

### Reconstruction of phylogeny based on signaling pathway genes

For each individual CDN signaling pathway gene, non-redundant amino acid sequences were aligned using MUSCLE, and a phylogenetic tree was constructed using the maximum likelihood (ML) method with the JTT model, incorporating 300 bootstrap replicates in MEGA 11. The resulting phylogenetic trees were visualized in iTOL, with identified domains from SMART marked accordingly (68). Domains visible in SMART were extracted using custom in-house scripts. For the *ataC* genes, some organisms displayed two or more gene representations overlapping with the *ataC* gene. As a result, we excluded the overlapping domains and focused solely on the *ataC* gene alongside non-overlapping genes. In the case of *M. goodii*, the *ataC* gene was not visible; however, it had a better E-value than the visible domain. Custom scripts were employed to represent the genes, and this representation was subsequently visualized in iTOL.

