## Supplementary material for "Exploring the evolutionary divergence of cyclic di-nucleotide signaling in diverse mycobacterial species": Supplemetal data

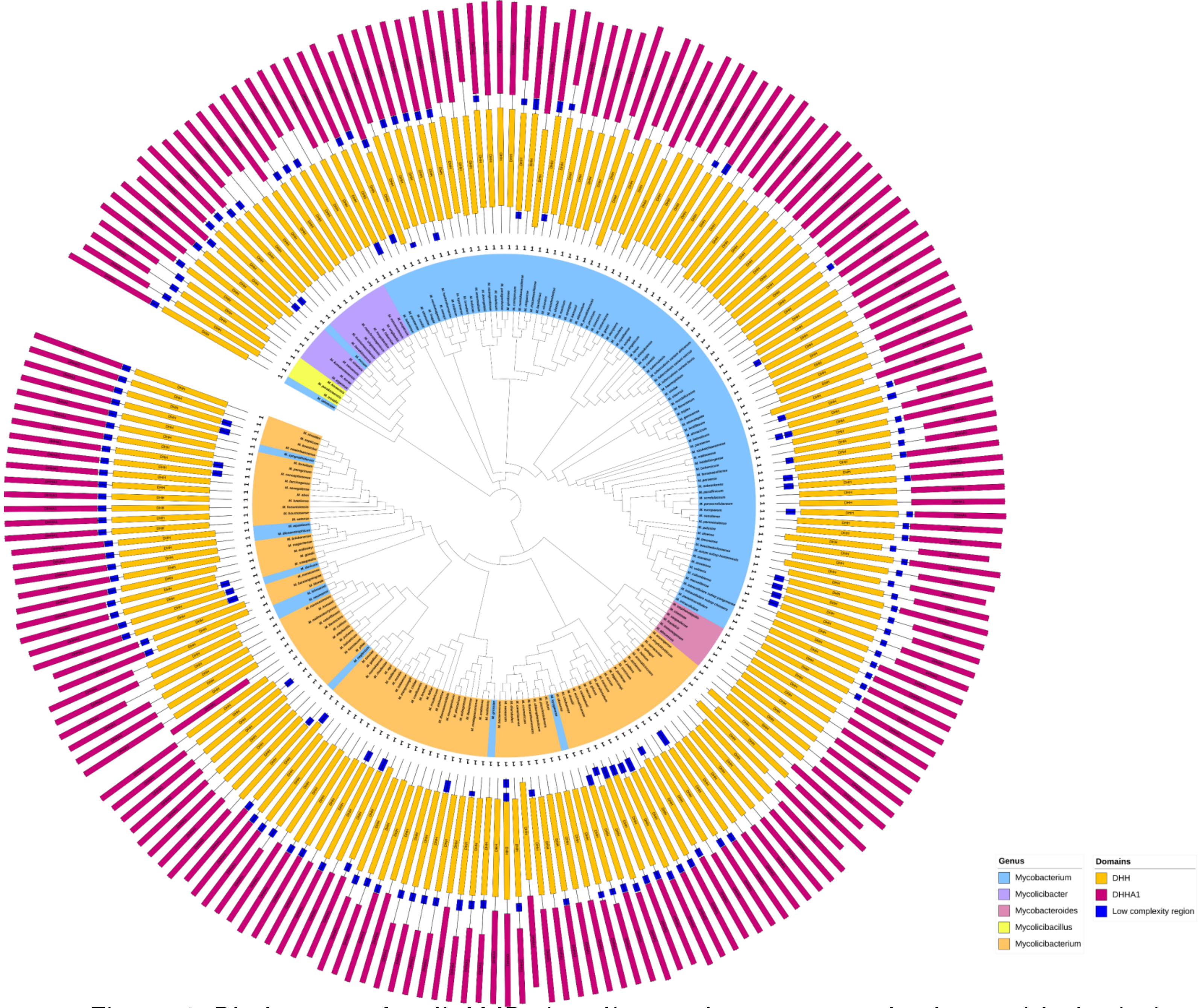


Supplementary Figure 1: Phylogeny of c-di-AMP signaling pathway gene *pde* along with depiction of distribution of functional domains

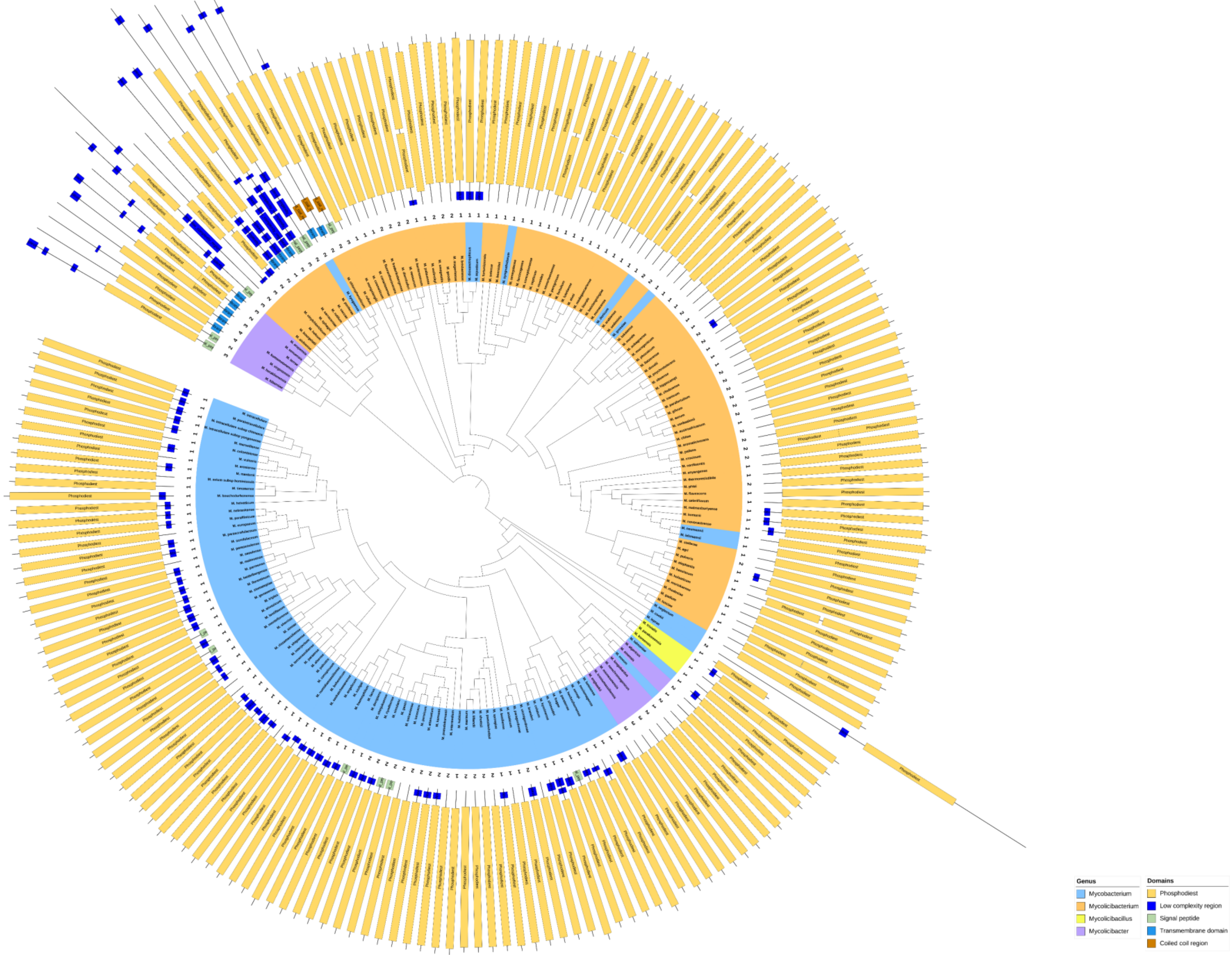


Supplementary Figure 2 : Phylogeny of c-di-AMP signaling pathway gene *ataC* along with depiction of distribution of functional domains

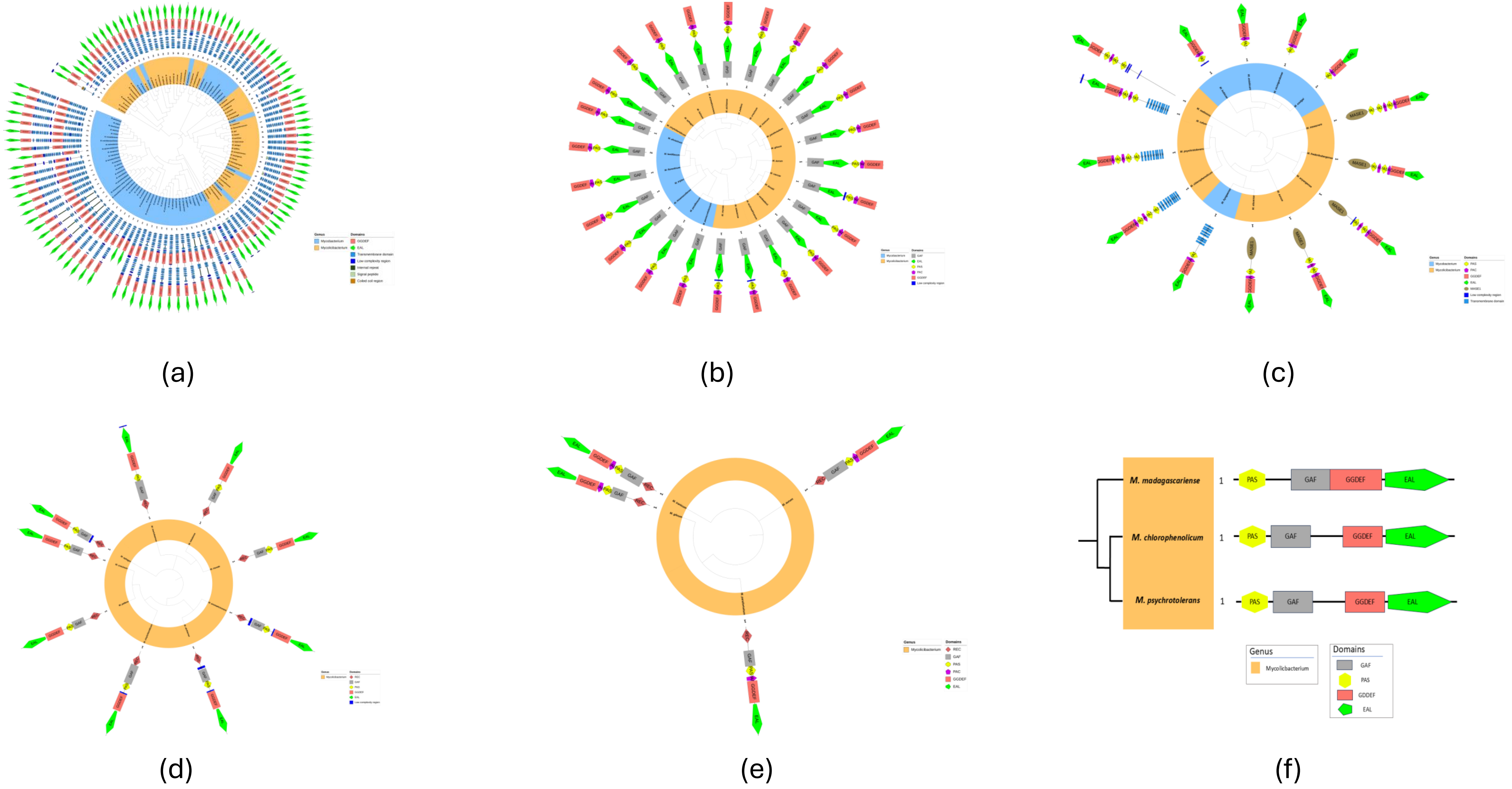


Supplementary Figure 3 : Phylogeny of c-di-GMP signaling pathway genes (a) *dcp*B, (b) *dcp*C, (c) *dcp*D, (e) *dcp*E, (f) *dcp*F, (g) *dcp*G along with depiction of distribution of functional domains

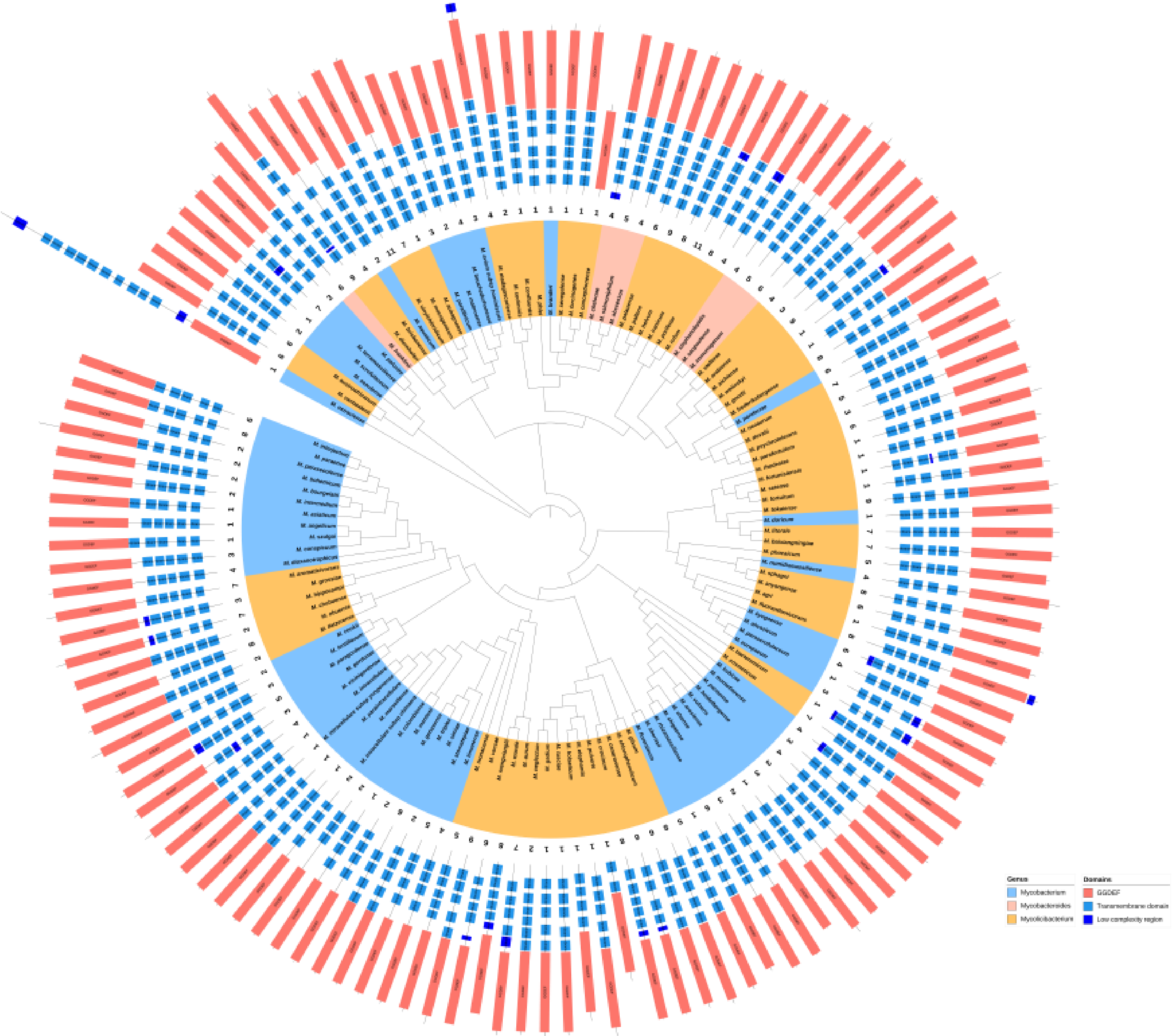


Supplementary Figure 4 : Phylogeny of c-di-GMP signaling pathway gene *dgc*A along with depiction of distribution of functional domains

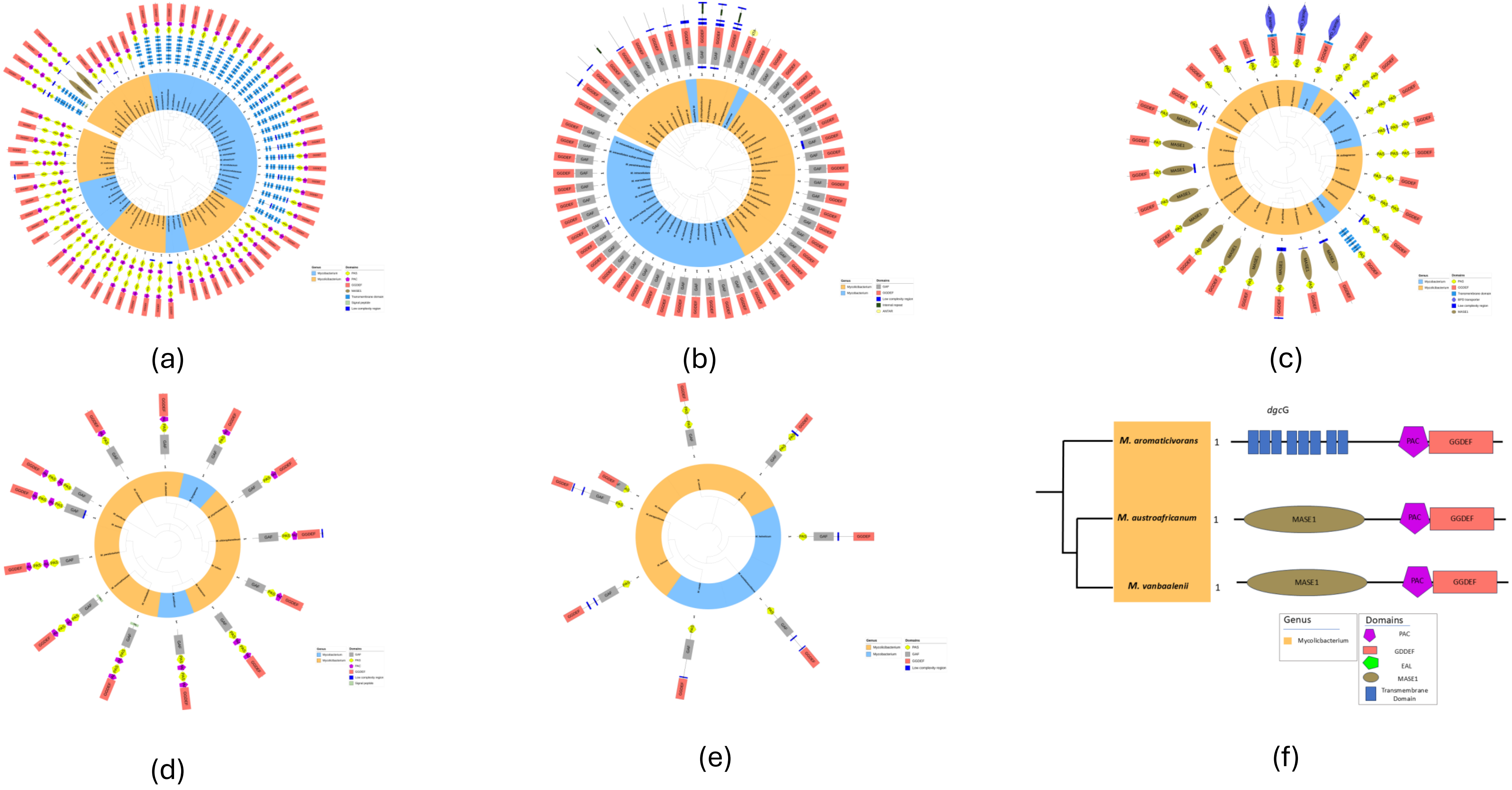


Supplementary Figure 5 : Phylogeny of c-di-GMP signaling pathway genes (a) *dgc*B, (b) *dgc*C, (c) *dgc*D, (d) *dgc*E, (e) *dgc*F, (f) *dgc*G along with depiction of distribution of functional domains

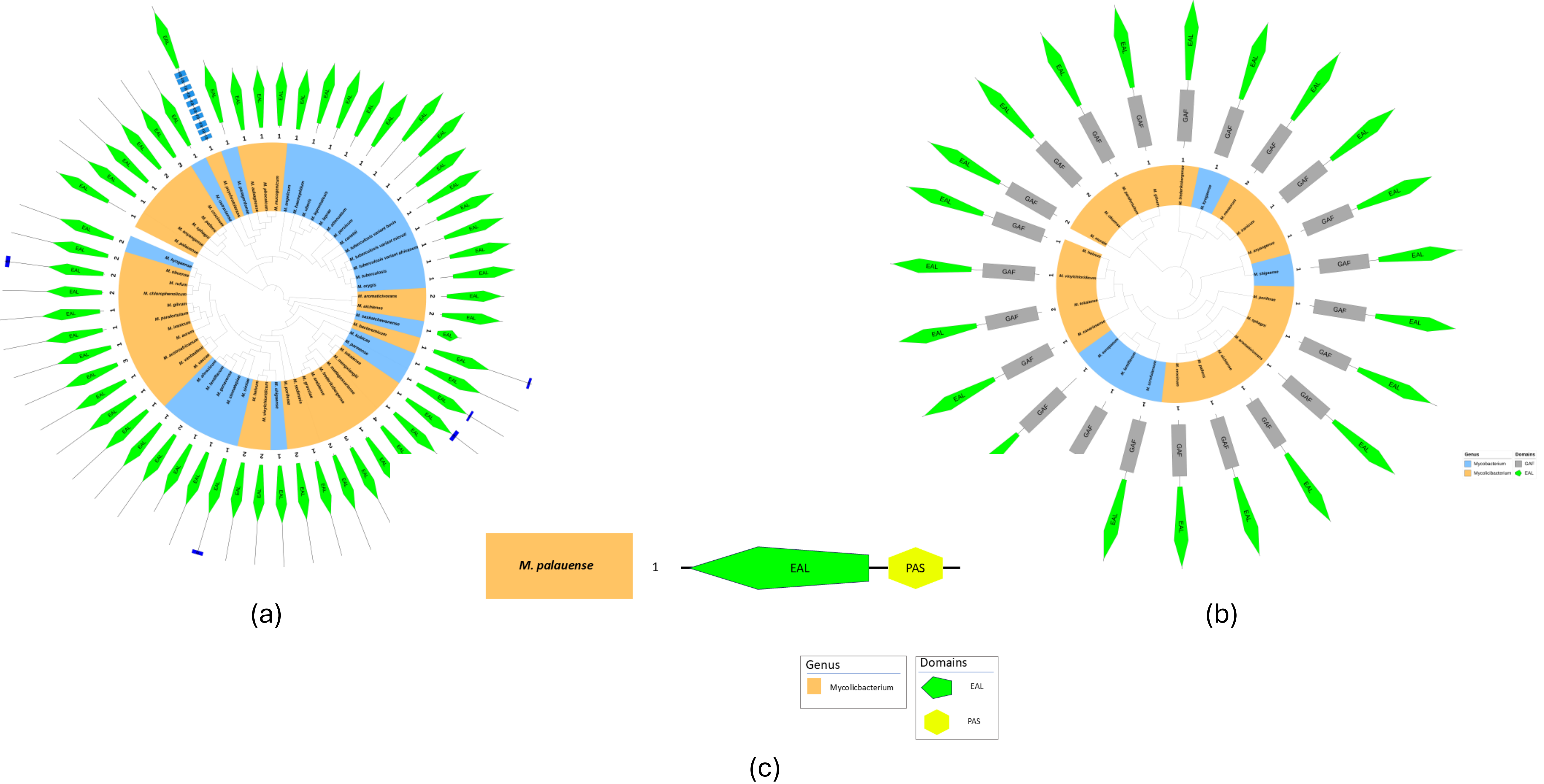


Supplementary Figure 6 : Phylogeny of c-di-GMP signaling pathway genes (a) *dgp*A, (b) *dgp*B, (c) *dgp*C along with depiction of distribution of functional domains

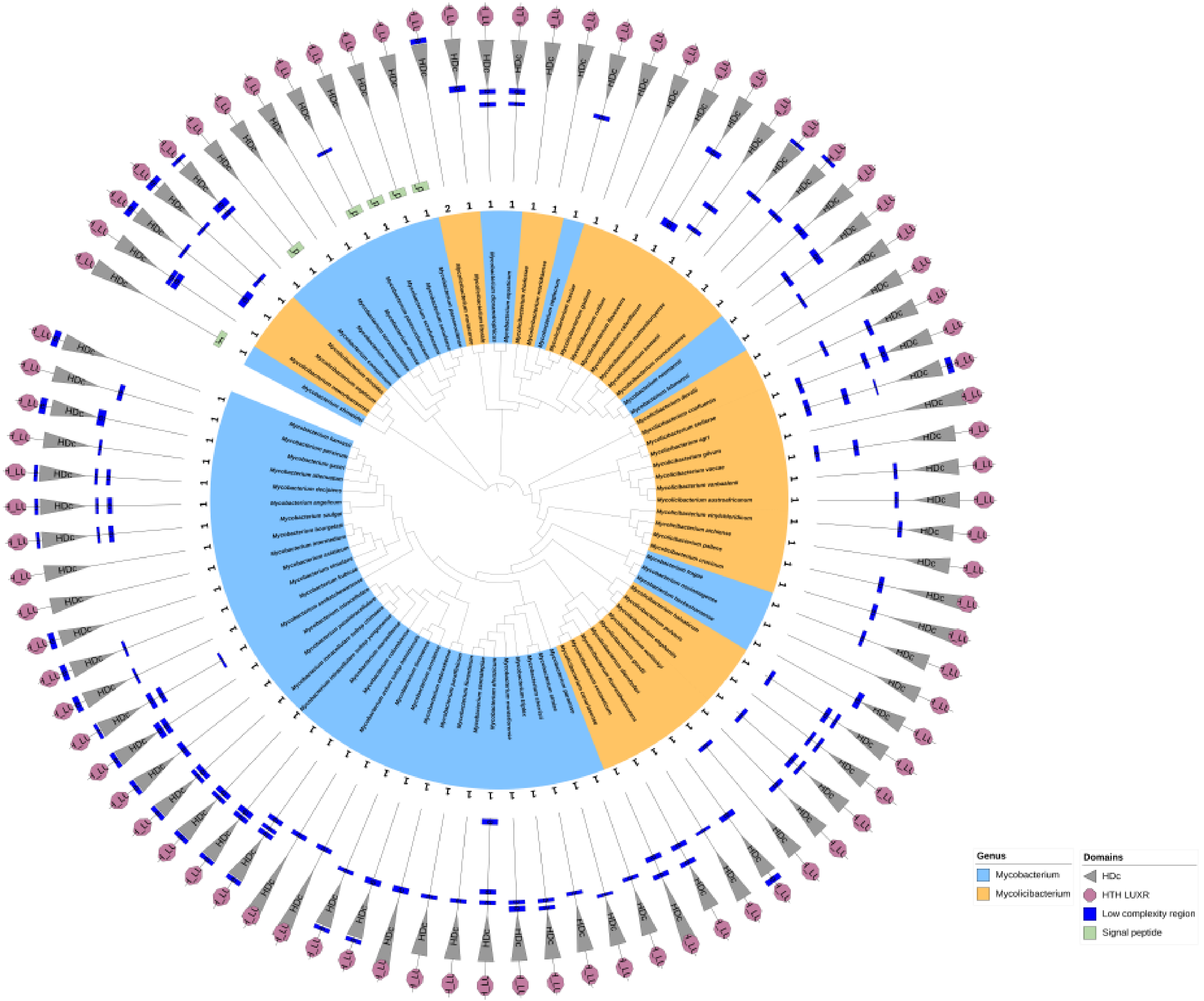


Supplementary Figure 7 : Phylogeny of c-di-GMP pathway genes containing the HD-GYP domain with mapped distribution of functional domains

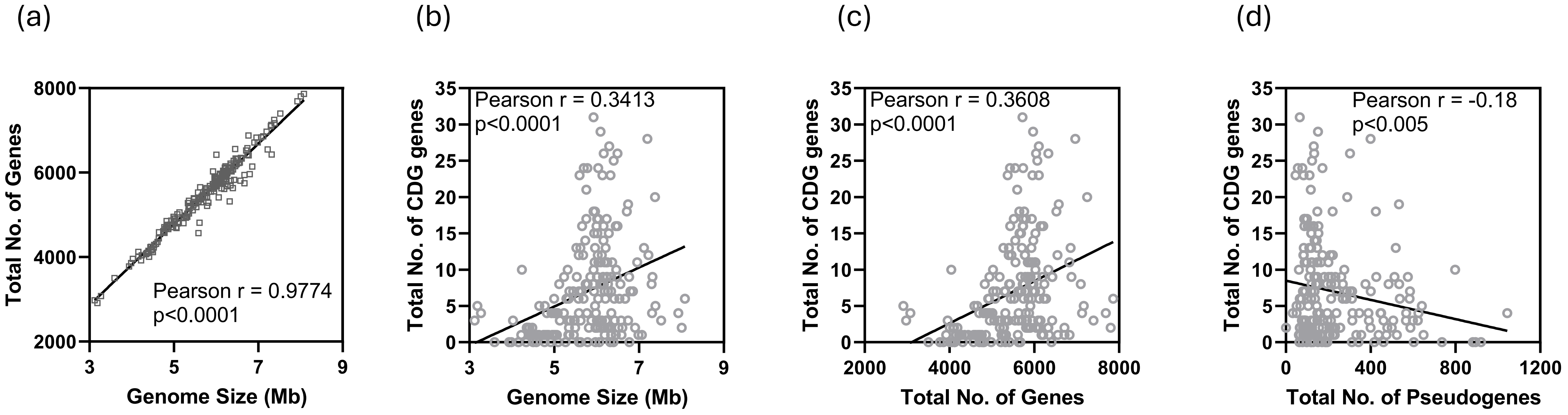


Supplementary Figure 8: Scatter plots depicting correlation between (a) genome size vs total number of genes present, (b) total number of c-di-GMP signaling pathways genes, (c) among all the organisms under study.
